# Photosynthetic induction upon transfer from low to high light is affected by leaf nitrogen content in tomato

**DOI:** 10.1101/2021.10.18.464921

**Authors:** Hu Sun, Yu-Qi Zhang, Shi-Bao Zhang, Wei Huang

## Abstract

The response of photosynthetic CO_2_ assimilation to changes of illumination affects plant growth and crop productivity under natural fluctuating light conditions. However, the effects of nitrogen (N) supply on photosynthetic physiology after transition from low to high light are seldom studied. To elucidate this, we measured gas exchange and chlorophyll fluorescence under fluctuating light in tomato (*Solanum lycopersicum*) seedlings grown with different N conditions. After transition from low to high light, the induction speeds of net CO_2_ assimilation (*A*_N_), stomatal conductance (*g*_s_) and mesophyll conductance (*g*_m_) delayed with the decline in leaf N content. The times to reach 90% of maximum *A*_N_, *g*_s_ and *g*_m_ were negatively correlated to leaf N content. This delayed photosynthetic induction in plants grown under low N concentration was mainly caused by the slow induction response of *g*_m_ rather than that of *g*_s_. Furthermore, the photosynthetic induction upon transfer from low to high light was hardly limited by photosynthetic electron flow. These results indicate that decreased leaf N content declines carbon gain under fluctuating light in tomato. Increasing the induction kinetics of *g*_m_ has the potential to enhance the carbon gain of field crops grown in infertile soil.

## Introduction

Plants capture light energy to produce chemical energy ATP and NADPH, which are used to drive nitrogen assimilation and the conversion of CO_2_ to sugar. Enhancing net CO_2_ assimilation rate (*A*_N_) is thought to be one of the most important targets for improving plant growth and crop productivity (Yamori *et al.* 2016a; Kromdijk *et al.* 2016; South, Cavanagh, Liu & Ort 2019). Many previous studies indicated that increasing *A*_N_ under constant high light can boost plant biomass (Kebeish *et al.* 2007; Timm *et al.* 2012, 2015). Recently, some studies reported that the response of *A*_N_ to increases of illumination significantly affects the carbon gain and thus influences plant growth (Slattery, Walker, Weber & Ort 2018; Adachi *et al.* 2019; Kimura, Hashimoto-Sugimoto, Iba, Terashima & Yamori 2020; Yamori, Kusumi, Iba & Terashima 2020; Zhang, Kaiser, Marcelis, Yang & Li 2020). Therefore, altering the photosynthetic performance under dynamic illumination is a promising way to improving photosynthesis under natural fluctuating light (FL) conditions.

Plants grown under high nitrogen (N) concentration usually have higher biomass than plants grown under low N concertation (Makino 2011). An important explanation for this is that leaf photosynthetic capacity is related to the leaf N content in many higher plants (Yamori, Nagai & Makino 2011; Li *et al.* 2020), since stromal enzymes and thylakoid proteins account for the majority of leaf N (Makino & Osmond 1991; Sudo, Makino & Mae 2003; Takashima, Hikosaka & Hirose 2004). Furthermore, stomatal conductance (*g*_s_) and mesophyll conductance (*g*_m_) under constant high light are also increased in plants grown under high N concentration, which speeds up CO_2_ diffusion from atmosphere to chloroplast carboxylation sites and thus favors the operation of *A*_N_ under constant high light (Yamori *et al.* 2011). However, little is known about the effects of leaf N content on non-steady-state photosynthetic performances under FL.

Under natural field conditions, light intensity exposed on leaf surface dynamically changes on timescales from milliseconds to hours (Pearcy 1990; Slattery *et al.* 2018). Furthermore, FL and N deficiency usually occurs concomitantly, but how FL and N deficiency interact to influence photosynthetic physiology in crop plants is poorly understood. After a sudden transitioning from low to high light, the gradual increase of *A*_N_ is termed “photosynthetic induction”. Recent studies indicated that the induction response of *A*_N_ was significantly affected by the induction speed of *g*_s_ (De Souza, Wang, Orr, Carmo-Silva & Long 2020; Kimura *et al.* 2020). Gene expression plays a crucial role in the induction response of *g*_s_ under FL. For example, the *slow anion channel-associated 1* (*slac1*), *open stomata 1* (*ost1*) and abscisic acid deficient *flacca* mutants, and the *proton ATPase translocation control 1* (*PATROL1*) overexpression line had faster stomatal opening responses than WT-types in *Arabidopsis thaliana*, rice and tomato (Kaiser, Morales, Harbinson, Heuvelink & Marcelis 2020; De Souza *et al.* 2020; Kimura *et al.* 2020; Yamori *et al.* 2020). Furthermore, the stomatal opening during photosynthetic induction can be affected by environment conditions such as target light intensity, magnitude of change, *g*_s_ at low light, the time of day and vapor pressure deficit (Kaiser *et al.* 2020; Sakoda *et al.* 2020; Eyland, van Wesemael, Lawson & Carpentier 2021). However, there have been few studies that examined the effect of leaf N content on the induction response of *g*_s_ after transition from low to high light (Li *et al.* 2020).

In addition to *g*_s_, *g*_m_ is a major factor affecting CO_2_ concentration in chloroplast, because *g*_m_ determines the CO_2_ diffusion from intercellular space into the chloroplast (Flexas *et al.* 2013; Carriquí *et al.* 2015). In general, *g*_m_ can be determined by structure across leaf profiles, genetic types, biochemical components and environmental conditions (Yamori *et al.* 2011; Xiong *et al.* 2015; Théroux-Rancourt & Gilbert 2017). Previous studies have highlighted that *g*_m_ is the most important limiting factor for *A*_N_ in many angiosperms (Peguero-Pina *et al.* 2017; Xiong, Douthe & Flexas 2018; Yang, Huang, Yang, Chang & Zhang 2018b). Short-term response of *g*_m_ to light intensity has been determined and found that it varies between plant species (Tazoe, Von Caemmerer, Badger & Evans 2009; Yamori, Evans & Von Caemmerer 2010a; Xiong *et al.* 2018; Yang, Hu & Huang 2020). However, the induction response of *g*_m_ after transition from low to high light is little known. The *g*_m_ level under constant light is also significantly affected by leaf N content (Yamori *et al.* 2011). Furthermore, the rapid responses of *g*_m_ to CO_2_ concentration and temperature were also affected by leaf N content (Xiong *et al.* 2015). However, no studies have elucidated the effect of leaf N content on induction response of *g*_m_ upon transfer from low to high light.

In this study, we aimed to characterize the effects of leaf N content on induction kinetics of *A*_N_, *g*_s_ and *g*_m_ after a sudden transition from low to high light. Gas exchange and chlorophyll fluorescence were measured in tomato plants grown under contrasting N concentrations. The dynamic limitations of *g*_s_, *g*_m_ and biochemical factors imposed on *A*_N_ were analyzed based on the biochemical model for C3 photosynthesis (Farquhar, von Caemmerer & Berry 1980). The effects of leaf N content on photosynthetic performances during photosynthetic induction were revealed.

## Materials and methods

### Plant materials and growth conditions

Tomato (*Solanum lycopersicum* cv. Hupishizi) plants were grown in a greenhouse with the light condition of 40% full sunlight. The day/night air temperatures were approximately 30/20°C, the relative air humidity was approximately 60%-70%, and the maximum light intensity exposed to leaves was approximately 800 μmol photons m^−2^ s^−1^. Plants were grown in 19-cm plastic pots with humus soil and the initial soil N content was 2.1 mg/g. Plants were fertilized with Peters Professional’s water solution (N:P:K = 15:4.8:24.1) or water as follows: high nitrogen (HN, 0.15 g N/plant every two days), middle nitrogen (MN, 0.05 g N/plant once a week) and low nitrogen (LN, 0 mM N/plant). 0.3% water solution were used for fertilization, and the nitrogen sources were 24% (NH_4_)_3_PO_4_, 65% KNO_3_ and 9.5% CH_4_N_2_O. To prevent any water stress, these plants were watered every day. After cultivation for one month, youngest fully developed leaves were used for measurements.

### Gas exchange and chlorophyll fluorescence measurements

An open gas exchange system (LI-6400XT; Li-Cor Biosciences, Lincoln, NE, USA) was used to simultaneously measure gas exchange and chlorophyll fluorescence. Measurements were performed at a leaf temperature of approximately 25°C, leaf-to-air vapour pressure deficit of 1.2-1.4 kpa, and flow rate of air through the system of 300 μmol/s. To measure photosynthetic induction after a short-term shadefleck, leaves were firstly adapted to a light intensity of 1500 μmol photons m^−2^ s^−1^ and air CO_2_ concentration of 400 μmol mol^−1^ for >20 min until *A*_N_ and g_s_ reached steady-state. Then, leaves were subjected to 5 min of low light (100 μmol photons m^−2^ s^−1^) followed by 30 min of high light (1500 μmol photons m^−2^ s^−1^), and gas exchange and chlorophyll fluorescence were logged every minute. iWUE was calculated as iWUE = *A*_N_/*g*_s_. The relative *A*_N_, *g*_s_ and *g*_m_ curves were obtained from the standardization against the maximum values after 30 min photosynthetic induction at high light. The time required to reach 90% of the maximum *A*_N_, *g*_s_ and *g*_m_ was estimated by the first time at which the relative values were higher than 90%. After photosynthetic induction measurement, the response of CO_2_ assimilation rate to incident intercellular CO_2_ concentration (*A*/*C*_i_) curves were measured by decreasing the CO_2_ concentration to a lower limit of 50 μmol mol^−1^ and then increasing stepwise to an upper limit of 1500 μmol mol^−1^. For each CO_2_ concentration, photosynthetic measurement was completed in 3 min. Using the *A*/*C*_i_ curves, the maximum rates of RuBP regeneration (*J*_max_) and carboxylation (*V*_cmax_) were calculated (Long & Bernacchi 2003).

The quantum yield of PSII photochemistry was calculated as Φ_PSII_ = (*F*_*m*_′ – *F*_*s*_)/*F*_*m*_′ (Genty, Briantais & Baker 1989), where *F*_*m*_′ and *F*_*s*_ represent the maximum and steady-state fluorescence after light adaptation, respectively (Baker 2004). The total electron transport rate through PSII (*J*_PSII_) was calculated as follows (Krall & Edwards 1992):

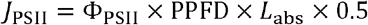

where PPFD is the photosynthetic photon flux density and leaf absorbance (*L*_abs_) is assumed to be 0.84. We applied the constant of 0.5 based on the assumption that photons were equally distributed between PSI and PSII.

### Estimation of mesophyll conductance and chloroplast CO_2_ concentration

Mesophyll conductance was calculated according to the following equation (Harley, Loreto, Di Marco & Sharkey 1992):

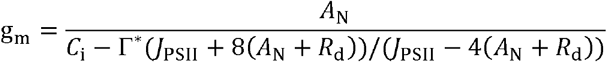

where *A*_N_ represents the net rate of CO_2_ assimilation; *C*_i_ is the intercellular CO_2_ concentration; Γ^*^ is the CO_2_ compensation point in the absence of daytime respiration (Yamori, Noguchi, Hikosaka & Terashima 2010b; von Caemmerer & Evans 2015), and we used a typical value of 40 μmol mol^−1^ in our current study (Xiong *et al.* 2018). Respiration rate in the dark (*R*_d_) was considered to be half of the dark-adapted mitochondrial respiration rate as measured after 10 min of dark adaptation (Carriquí *et al.* 2015).

Based on the estimated *g*_m_, we then calculated the chloroplast CO_2_ concentration (*C*_c_) according to the following equation (Long & Bernacchi 2003; Warren & Dreyer 2006):

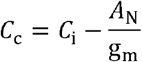

### Quantitative limitation analysis of *A*_N_

Relative photosynthetic limitations were assessed as follows (Grassi & Magnani 2005):

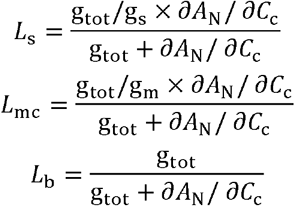

where *L*_s_, *L*_mc_, and *L*_b_ represent the relative limitations of stomatal conductance, mesophyll conductance, and biochemical capacity, respectively, in setting the observed value of *A*_N_. *g*_tot_ is the total conductance of CO_2_ between the leaf surface and sites of RuBP carboxylation (calculated as 1/*g*_tot_ = 1/*g*_s_ + 1/*g*_m_).

### SPAD index and nitrogen content measurements

A handy chlorophyll meter (SPAD-502 Plus; Minolta, Tokyo, Japan) was used to non-destructively measure the SPAD index (relative content of chlorophyll per unit leaf area) of leaves used for photosynthetic measurements. Thereafter, leaf area was measured using a LI-3000A portable leaf area meter (Li-Cor, Lincoln, NE, USA). After leaf material was dried at 80°C for 48 hours, dry weight was measured and leaf N content was determined with a Vario MICRO Cube Elemental Analyzer (Elementar Analysensysteme GmbH, Langenselbold, Germany) (Sakowska *et al.* 2018).

## Results

### Effect of leaf N content on steady-state physiological characteristics under high light

The leaf N content in LN-, MN- and HN-plants were 0.42 ± 0.03, 0.71 ± 0.3 and 1.2 ± 0.07 g m^−2^, respectively (Table 1). The HN-plants displayed the highest relative chlorophyll content, measured by SPAD value, followed by MN- and LN-plants. After 30 min light adaptation at 1500 μmol photons m^−2^ s^−1^ and 400 μmol mol^−1^ CO_2_ concentration, HN-plants had the highest net CO_2_ assimilation rate (*A*_N_), stomatal conductance (*g*_s_), mesophyll conductance (*g*_m_) and electron transport rate (ETR). Therefore, the steady-state photosynthetic capacities were significantly affected by leaf N content. Furthermore, HN-, MN- and LN-plants showed slight difference in *g*_s_ but significant difference in *g*_m_, indicating that *g*_m_ is more responsive to leaf N content than *g*_s_ in tomato.

**Table 1.**
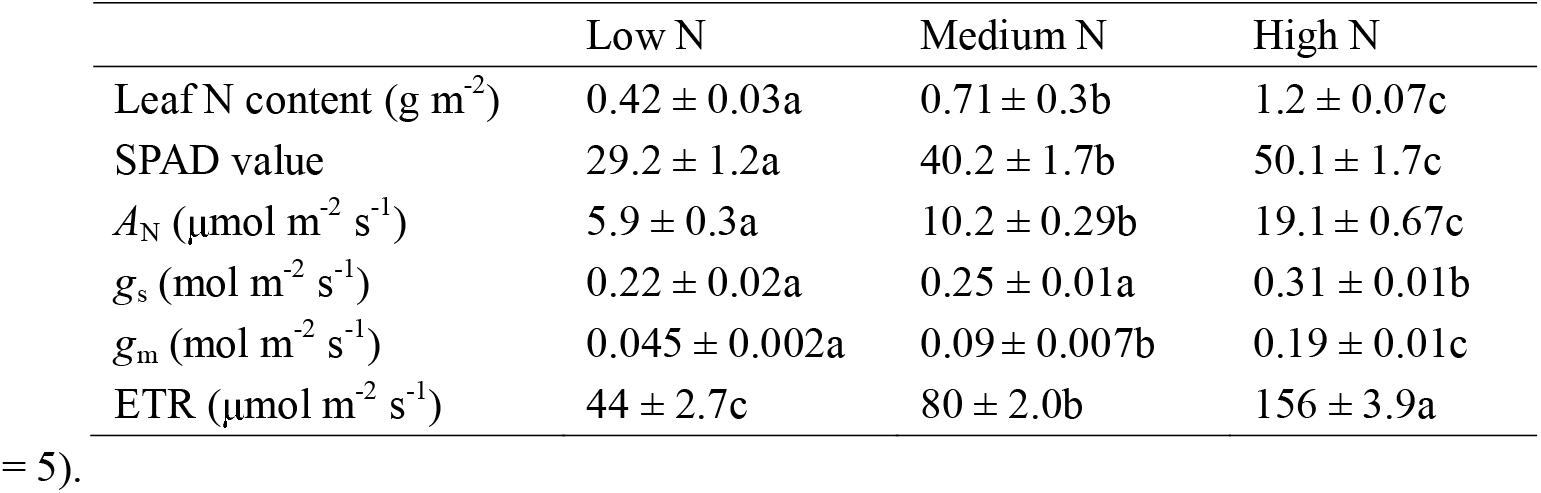
Physiological characteristics of leaves from plants grown under three different nutrient concentrations (low, medium and high nitrogen). All parameters were measured at 1500 μmol photons m^−2^ s^−1^ and 400 μmol mol^−1^ CO2 concentration. Values are means ± SE (n = 5).

### Effects of leaf N content on photosynthetic induction upon transfer from low to high light

During this photosynthetic induction after 5 min of shadefleck, HN-plants showed the highest induction speeds of *A*_N_, *g*_s_ and *g*_m_, followed by MN- and LN-plants (Fig. 1). The time required to reach 90% of the maximum *A*_N_ (*t*_90AN_) significantly increased with the decrease in leaf N content (Fig. 1G). The time required to reach 90% of the maximum *g*_s_ and *g*_m_ (*t*_90gs_ and *t*_90gm_, respectively) were significantly shorter in HN-plants than MN- and LN-plants, whereas *t*_90gs_ and *t*_90gm_ did not differ significantly between MN- and LN-plants (Fig. 1G). Interestingly, *t*_90gm_ was lower than *t*_90gs_ in all plants. The higher *t*_90gs_ and *t*_90AN_ in MN- and LN-plants was partially related to the relatively lower initial *g*_s_ prior to light change (Fig. S1). Within the first 15 minutes after transition from low to high light, all plants showed similar intrinsic water use efficiency (iWUE) (Fig. S2). However, during prolonged photosynthetic induction, HN-plants displayed much higher iWUE than MN- and LN-plants (Fig. S2). Further analysis found that leaf N content was negatively correlated to *t*_90AN_, *t*_90gs_ and *t*_90gm_ (Fig. 2). Therefore, leaf N content plays a crucial role in affecting the induction responses of *A*_N_, *g*_s_ and *g*_m_ after transition from low to high light. The comparative extent of the reductions of *t*_90AN_ was more correlated to *t*_90gm_ than *t*_90gs_ (Fig. 3A). Furthermore, the change in *A*_N_ during photosynthetic induction was more related to *g*_m_ than *g*_s_ (Fig. 3B&C). These results suggest that, upon transfer from low to high light, *g*_m_ plays a more important role in determining the induction response of *A*_N_ than *g*_s_.

**Figure 1.**
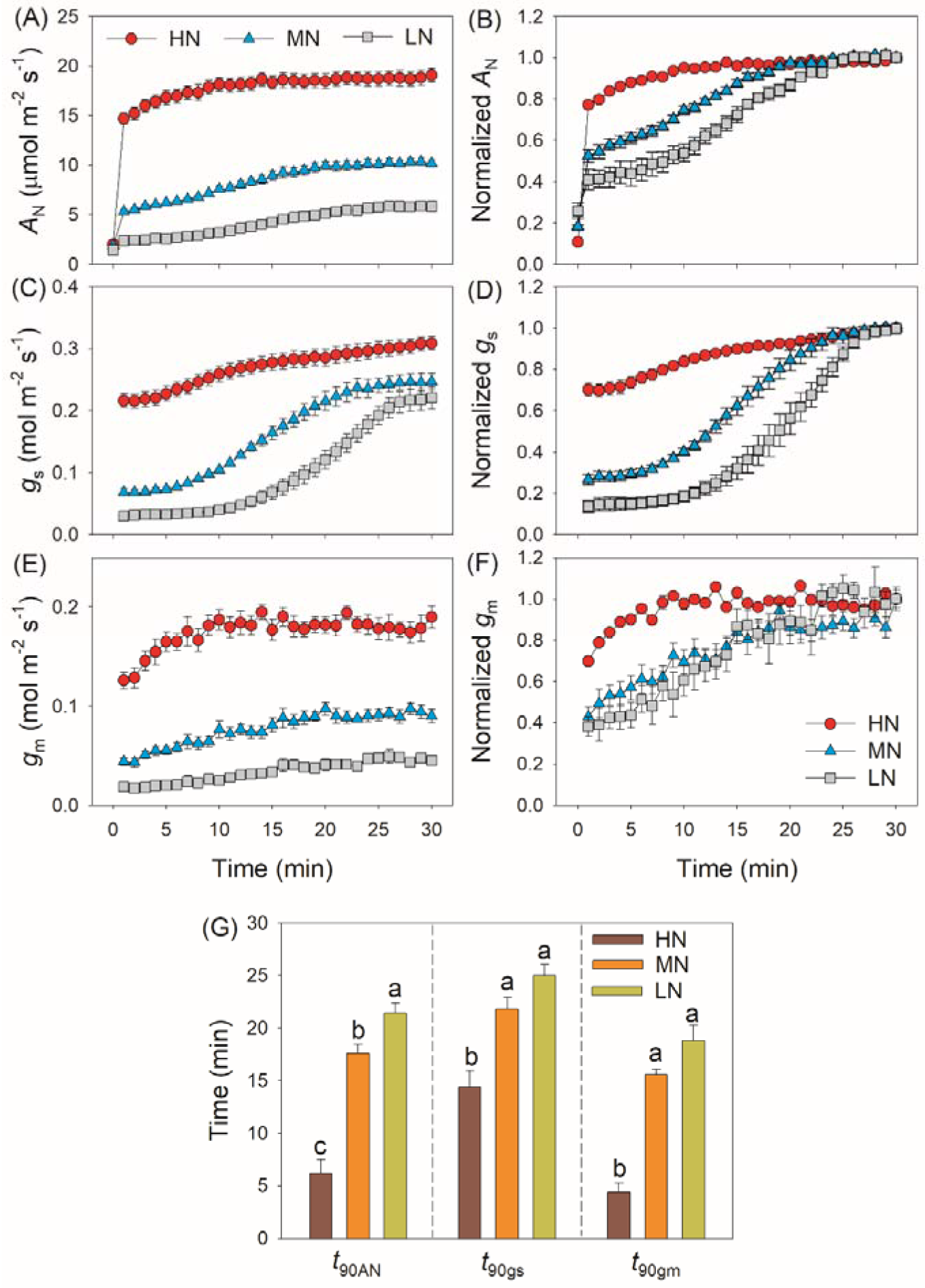
Induction response of net CO_2_ assimilation rate (*A*_N_), stomatal conductance (*g*_s_) and mesophyll conductance (*g*_m_), and the time required to reach 90% of the maximum values of *A*_N_, *g*_s_ and *g*_m_ (*t*_90AN_, *t*_90gs_, *t*_90gm_) after transition from 50 to 1500 μmol photons m^−2^ s^−1^. *A*_N_, *g*_s_ and *g*_m_ were measured every 1 min. Values are means ± SE (n = 5). Different letters indicate significant differences among different treatments. The relative *A*_N_, *g*_s_ and *g*_m_ curves were obtained from the standardization against the maximum values after 30 min photosynthetic induction at high light. HN, MN and LN represent tomato plants grown under high, medium and low N concentrations, respectively.

**Figure 2.**
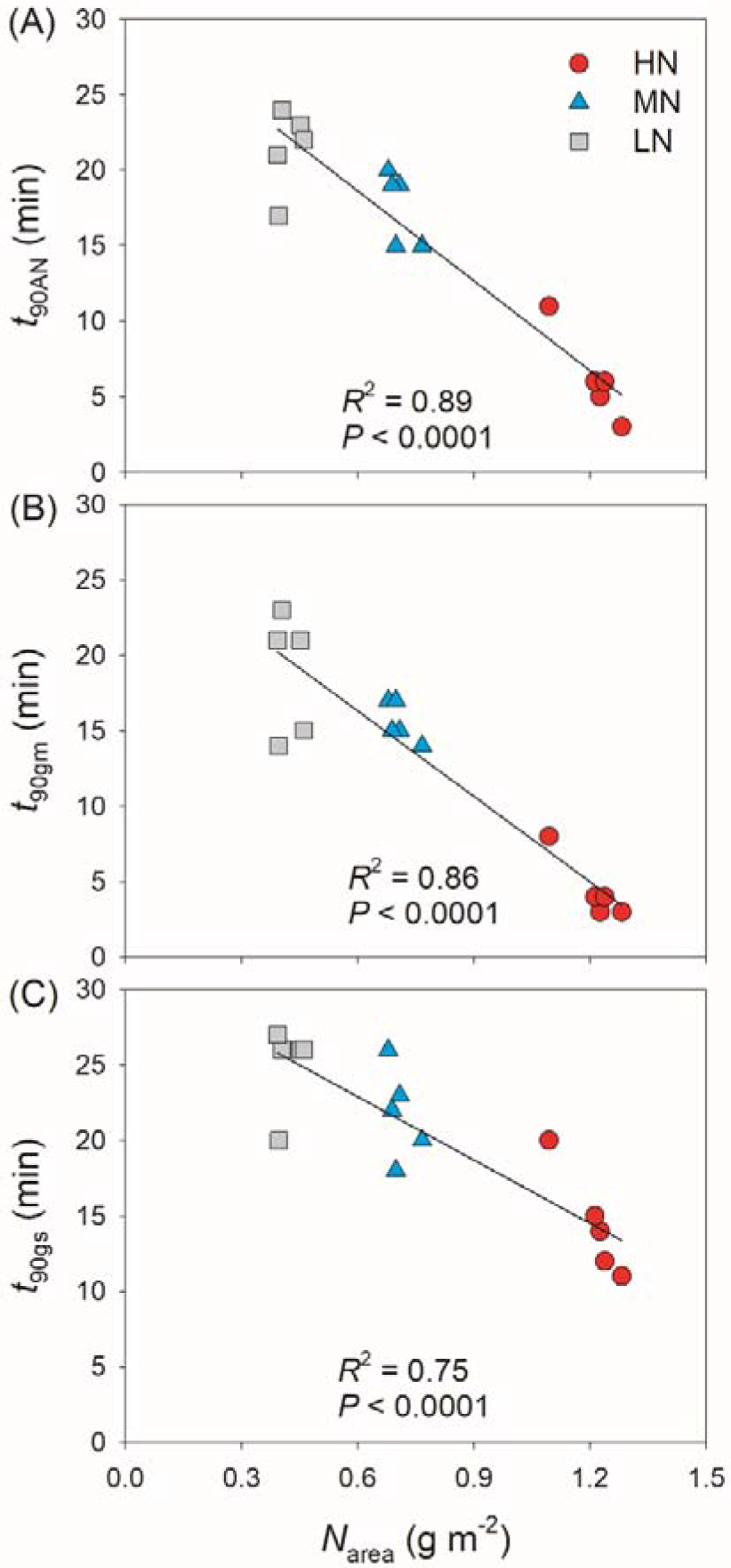
Effects of leaf N content on the time required to reach 90% of the maximum values of *A*_N_, *g*_s_ and *g*_m_ (*t*_90AN_, *t*_90gs_, *t*_90gm_) after transition from 50 to 1500 μmol photons m^−2^ s^−1^. HN, MN and LN represent tomato plants grown under high, medium and low N concentrations, respectively.

**Figure 3.**
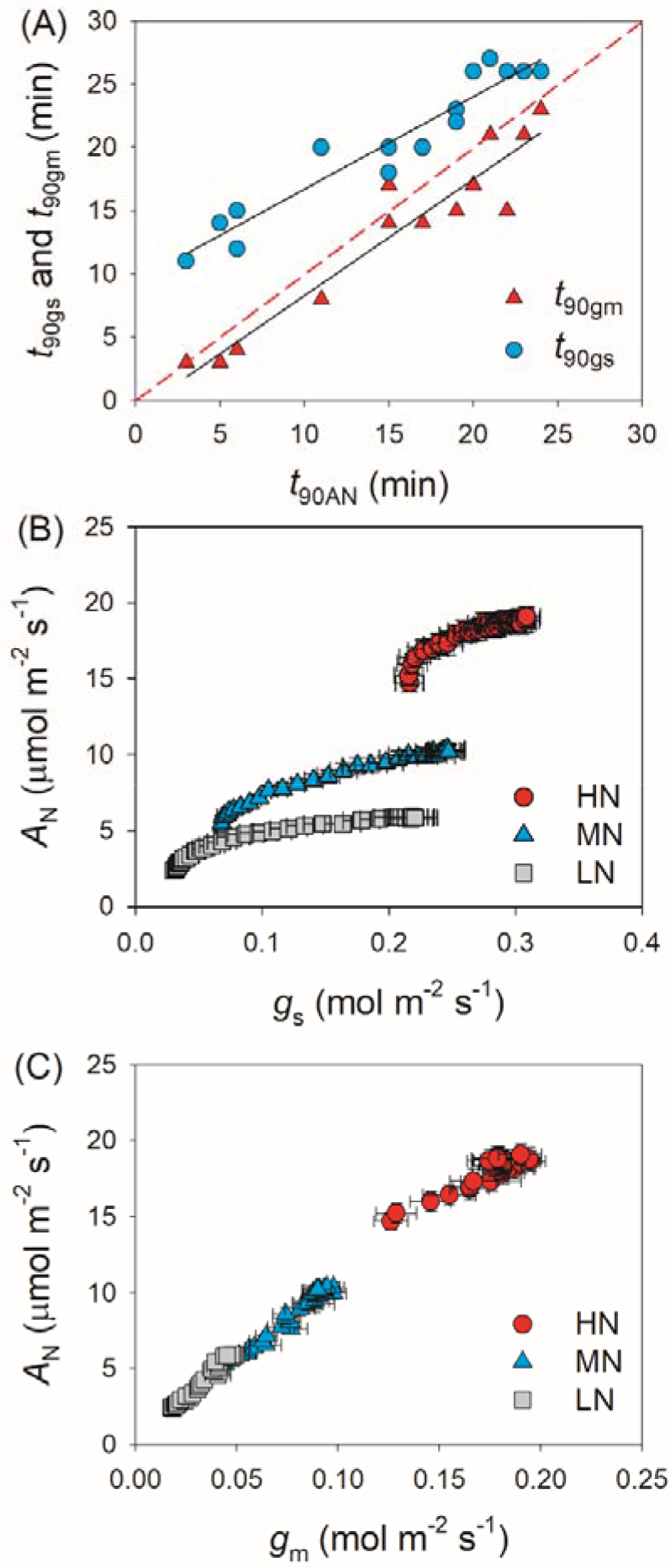
(**A**) Relationships between *t*_90AN_, *t*_90gs_ and *t*_90gm_ after transition from 50 to 1500 μmol photons m^−2^ s^−1^. (**B** and **C**) Relationships between *g*_s_, *g*_m_ and *A*_N_ after transition from 50 to 1500 μmol photons m^−2^ s^−1^. Values are means ± SE (n = 5). HN, MN and LN represent tomato plants grown under high, medium and low N concentrations, respectively.

### Effects of leaf N content on intercellular and chloroplast CO_2_ concentrations upon transfer from low to high light

We calculated the response kinetics of intercellular (*C*_i_) and chloroplast CO_2_ concentration (*C*_c_) using *A*_N_, *g*_s_ and *g*_m_. After transitioning from low to high light, *C*_i_ and *C*_c_ gradually increased in all plants (Fig. 4). HN-plants had the lowest values of *C*_i_ and *C*_c_ after photosynthetic sufficient photosynthetic induction. The change in *A*_N_ during photosynthetic induction was tightly and positively correlated to *C*_c_ in all plants, suggesting the importance of *C*_c_ in determining *A*_N_. Because *C*_c_ can be affected by *g*_s_ and *g*_m_, we analyzed the relationships between *C*_c_, *g*_s_ and *g*_m_. Compared with *g*_s_, a smaller change in *g*_m_ could result in a larger change in *C*_c_ (Fig. 5), suggesting that the change of *C*_c_ upon transfer from low to high light was more determined by *g*_m_ than *g*_s_.

**Figure 4.**
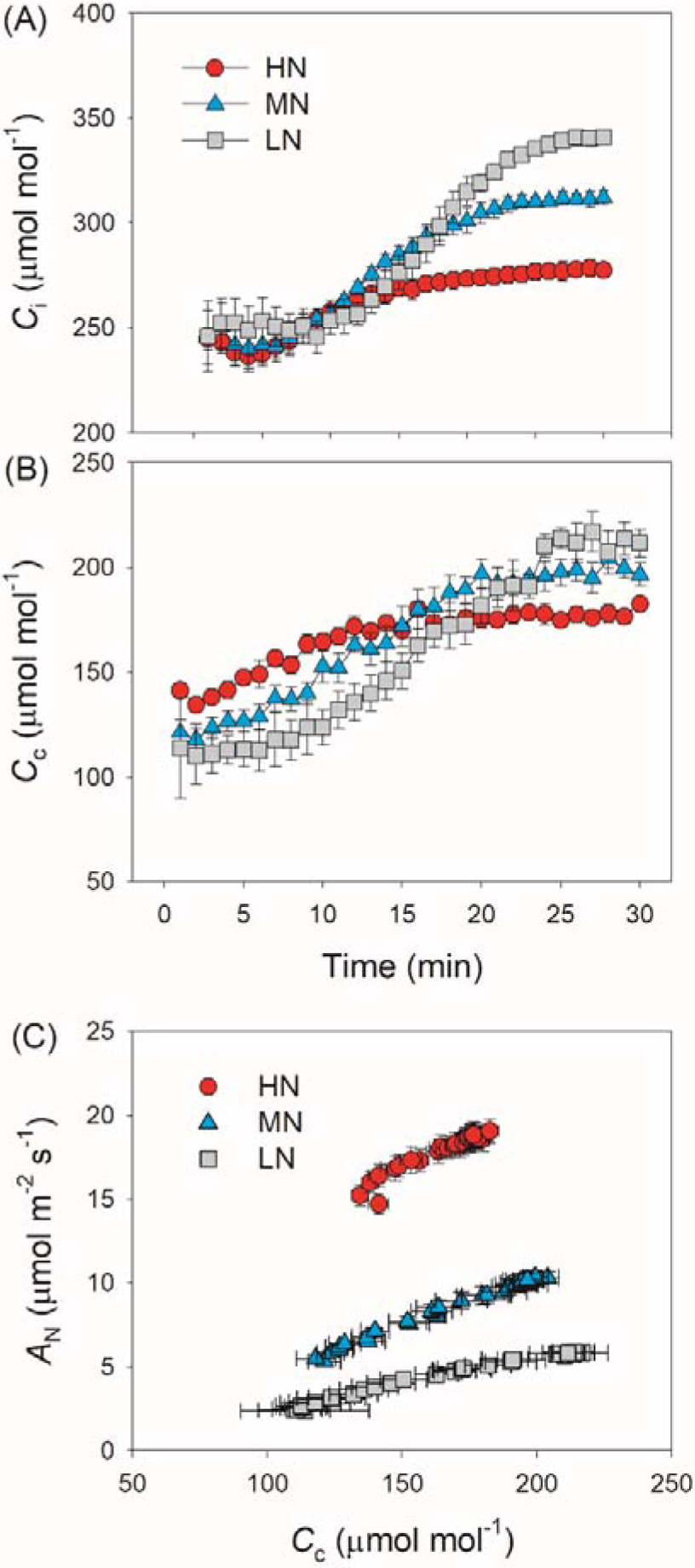
(**A** and **B**) Response of intercellular CO_2_ concentration (*C*_i_) and chloroplast CO_2_ concentration (*C*_c_) after transition from 50 to 1500 μmol photons m^−2^ s^−1^. (**C**) Relationship between *C*_c_ and *A*_N_ after transition from 50 to 1500 μmol photons m^−2^ s^−1^. Values are means ± SE (n = 5). HN, MN and LN represent tomato plants grown under high, medium and low N concentrations, respectively.

**Figure 5.**
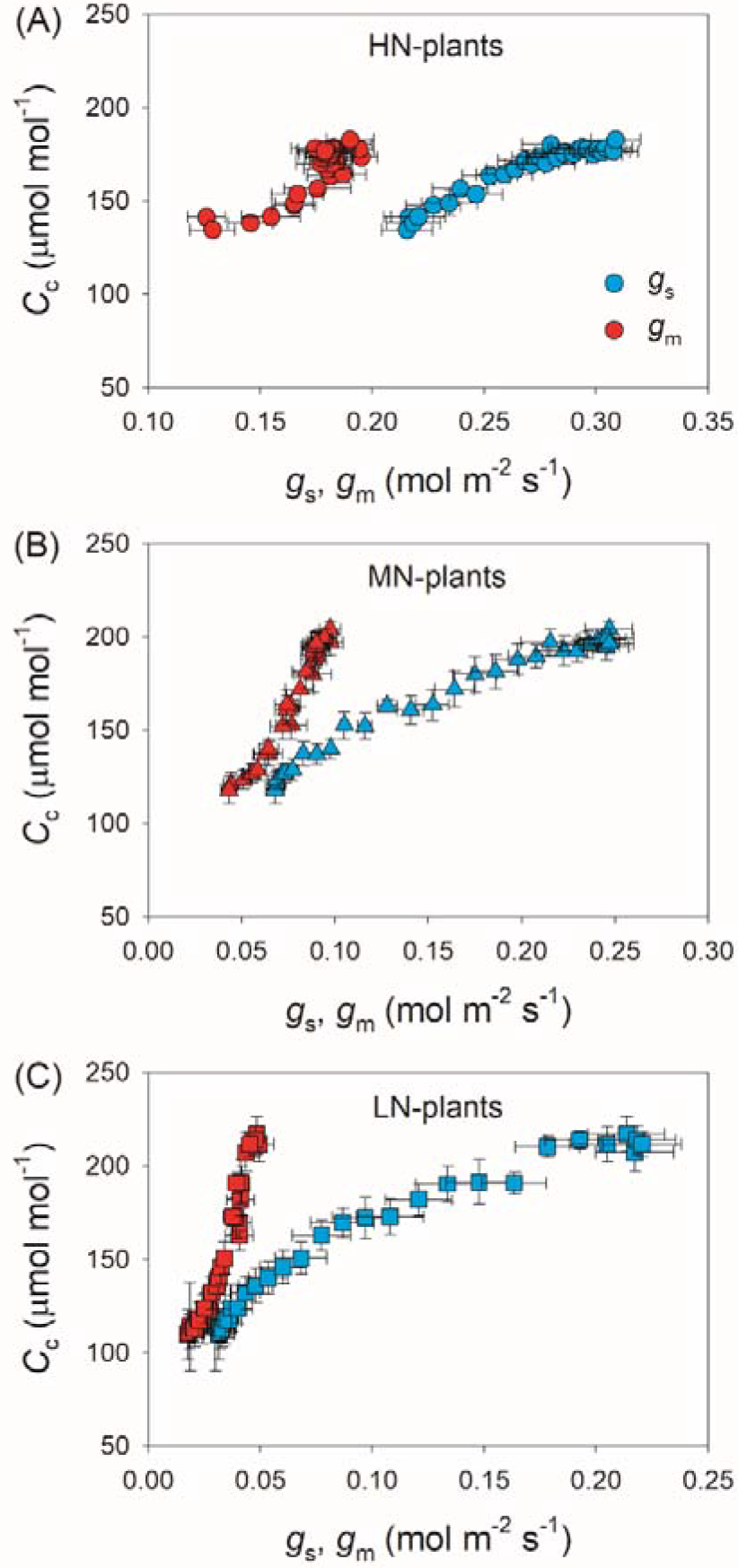
Relationships between *g*_s_, *g*_m_ and *C*_c_ after transition from 50 to 1500 μmol photons m^−2^ s^−1^. Values are means ± SE (n = 5). HN, MN and LN represent tomato plants grown under high, medium and low N concentrations, respectively.

### Effects of leaf N content on relative limitations of photosynthesis upon transfer from low to high light

After transition from low to high light, the limitations of photosynthesis by *g*_s_ (*L*_gs_), *g*_m_ (*L*_gm_) and biochemical factors (*L*_b_) changed slightly in HN-plants (Fig. 6). In MN- and LN-plants, *L*_gs_ gradually decreased over time. Within the first 15 min, *L*_gs_ was lower in HN-plants than MN- and LN-plants. However, the LN-plants had the lowest *L*_gs_ after sufficient photosynthetic induction. *L*_gm_ was also maintained stable during whole photosynthetic induction in MN- and LN-plants, but *L*_b_ gradually increased from 0.3 to 0.5 in them. Therefore, leaf N content could affect the kinetics of relative limitations of photosynthesis during photosynthetic induction after transfer from low to high light. To explore whether the induction of *A*_N_ is limited by photosynthetic electron transport, we estimated the dynamic change of electron transport rate (ETR). Upon a sudden increase in illumination, ETR rapidly increased and the ETR/(*A*_N_ + *R*_d_) ratio first increased and then gradually decreased in all plants (Fig. 7). These results indicated that the activation speed of ETR was much faster than that of *A*_N_. Therefore, during photosynthetic induction the limitation of ETR imposed to *A*_N_ was negligible in all samples.

**Figure 6.**
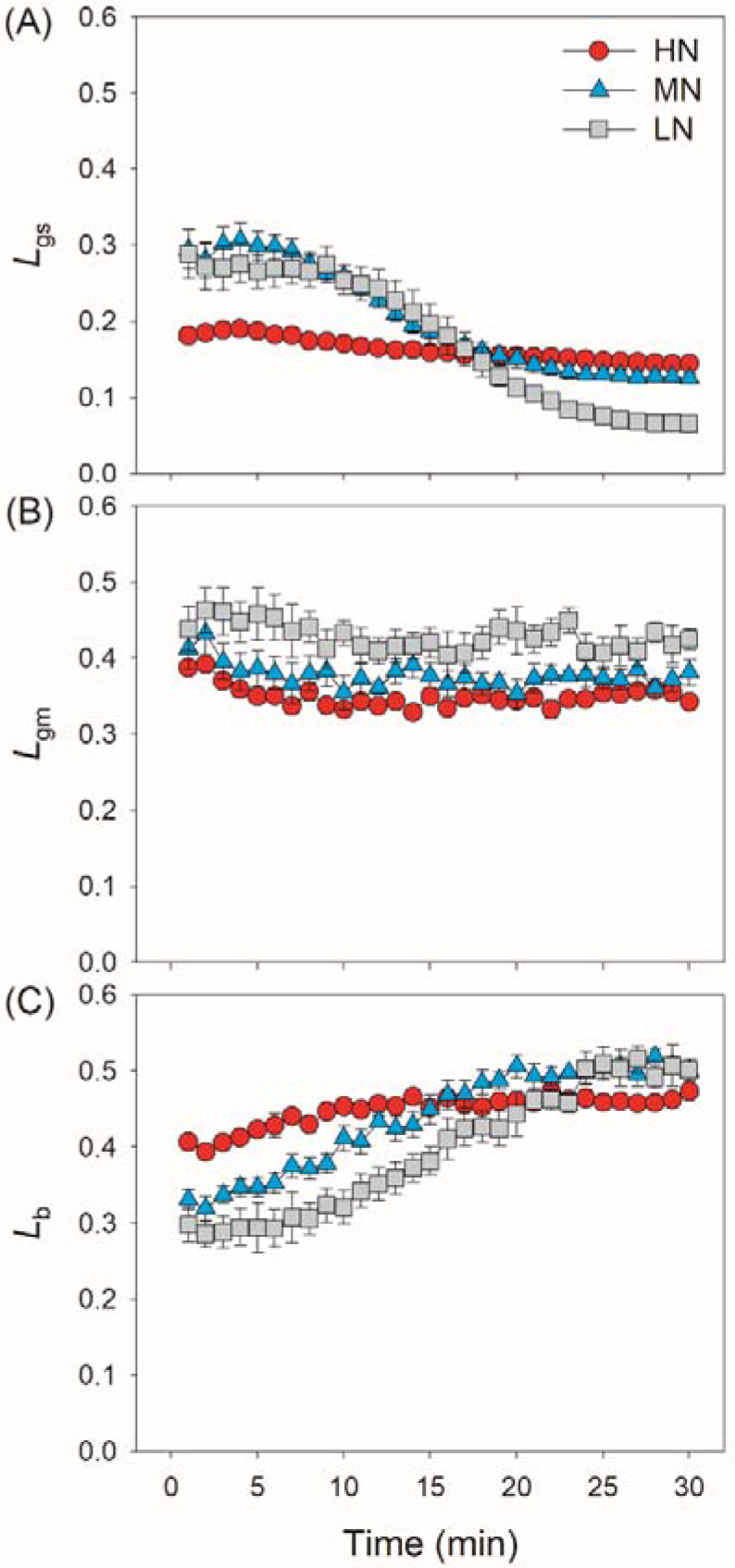
Quantitative analysis of the relative limitations of *g*_s_, *g*_m_ and biochemical factors imposed to photosynthesis after transition from 50 to 1500 μmol photons m^−2^ s^−1^. Values are means ± SE (n = 5). HN, MN and LN represent tomato plants grown under high, medium and low N concentrations, respectively.

**Figure 7.**
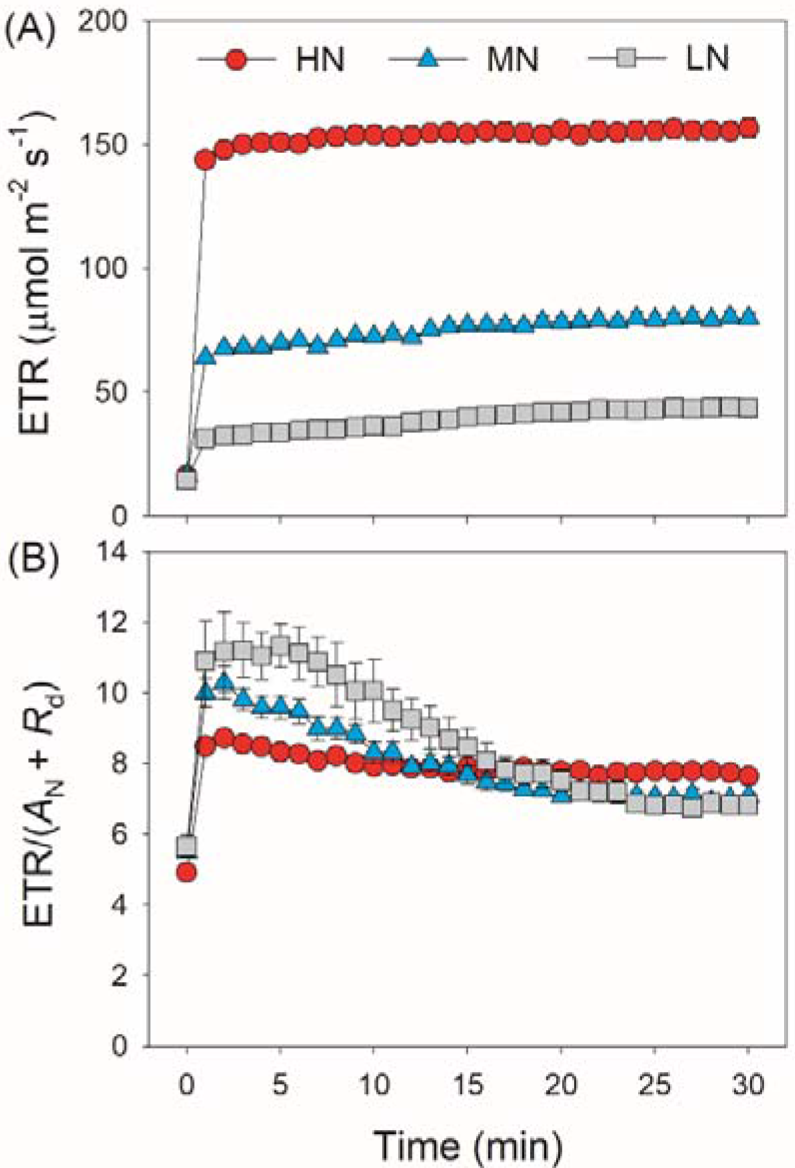
Response of electron transport rate (ETR) and the ratio of ETR to (*A*_N_ + *R*_d_) after transition from 50 to 1500 μmol photons m^−2^ s^−1^. Values are means ± SE (n = 5). HN, MN and LN represent tomato plants grown under high, medium and low N concentrations, respectively.

## Discussion

Leaf N content plays an important role in determining photosynthesis, plant growth and crop productivity (Makino 2011). Under natural field conditions, FL and N deficiency usually occurs concomitantly. However, it is unknown how FL and N deficiency interact to influence photosynthetic physiology in crop plants. In this study, we here for the first time examined the effects of leaf N content on photosynthetic induction after transition from low to high light in tomato. We found that leaf N content significantly affected the induction responses of *g*_s_ and *g*_m_ and thus affected induction kinetics of *A*_N_. However, the activation speed of photosynthetic electron flow was not influenced by leaf N content. Therefore, the effect of leaf N content on photosynthetic induction was more attributed to the induction kinetics of diffusional conductance rather than the activation speed of electron transport.

In addition to steady-state photosynthetic capacity under high light, the photosynthetic responses to the changes in illumination significantly affect the carbon gain and plant biomass (Adachi *et al.* 2019; Kimura *et al.* 2020; Zhang *et al.* 2020). Many previous studies have documented that leaf N content influences the steady-state photosynthetic performances under high light (Evans & Terashima 1988; Makino & Osmond 1991), but little is known about the influence of leaf N content on photosynthetic induction under FL conditions. Similar to previous studies, the maximum steady-state *A*_N_ under high light significantly declined with the decrease in leaf N content (Table 1). Moreover, we here found that, after transition from low to high light, the HN-plants showed much faster induction response of *A*_N_ than MN- and LN-plants (Fig. 1). The time required to reach 90% of the steady-state of photosynthesis (*t*_90AN_) was negatively correlated to leaf N content (Fig. 2). Therefore, leaf N content significantly affect the photosynthetic induction after transition from low to high light in tomato. This finding is similar to the photosynthetic induction of dark-adapted leaves among canola genotypes (*Brassica napus* L.) (Liu, Zhang, Estavillo, Luo & Hu 2021), but was inconsistent with the phenomenon in soybean (Li *et al.* 2020). In soybean, the induction rate of *A*_N_ under high light after shading for 5 min was very fast (Pearcy, Krall & Sassenrath-Cole 1996; Li *et al.* 2020). Furthermore, this fast photosynthetic induction in soybean was not affected by leaf N content (Li *et al.* 2020). Therefore, the effect of leaf N content on fast photosynthetic induction following shadefleck depends on the species and on growth conditions. In MN- and LN-plants of tomato, the delayed induction of *A*_N_ caused a larger loss of carbon gain under FL. This finding provides insight into why plants grown under low N concentrations display reduction of plant biomass under natural field FL conditions.

After transition from low to high light, the time to reach the maximum *C*_c_ was less in HN-plants than MN- and LN-plants (Fig. 4). Furthermore, tight and positive relationships were found between *C*_c_ and *A*_N_ in all plants (Fig. 4). These results suggested that the induction response of *A*_N_ was largely determined by the change of CO_2_ concentration in the site of RuBP carboxylation. The value of *C*_c_ in a given leaf is largely affected by CO_2_ diffusional conductance, including *g*_s_ and *g*_m_ (Sagardoy *et al.* 2010; Carriquí *et al.* 2015; Yang *et al.* 2018b). However, it is unclear whether the photosynthetic induction of *A*_N_ upon transfer from low to high light is more determined by the induction response of *g*_s_ or *g*_m_. We found that the induction responses of *g*_s_ and *g*_m_ were largely delayed in MN- and LN-plants than HN-plants (Fig. 1), and the induction rates of *g*_s_ and *g*_m_ were negatively correlated to leaf N content (Fig. 2). Furthermore, the change of *C*_c_ during photosynthetic induction was more related to *g*_m_ rather than *g*_s_ (Fig. 5), pointing out the important role of *g*_m_ response in determining *C*_c_ upon transfer from low to high light. Therefore, the delayed photosynthetic induction of *A*_N_ in plants grown under low N concentrations was more attributed to the slower induction response of *g*_m_ than *g*_s_.

In HN-plants of tomato, photosynthetic limitations by *g*_s_, *g*_m_ and biochemical factors changed slightly upon transfer from low to high light. Meanwhile, *g*_s_ imposed to the smallest limitation to *A*_N_, owing to the high levels of *g*_s_ (Fig. 6). Therefore, improving the induction response of *g*_s_ might have a minor factor for improving photosynthesis under FL in HN-plants of tomato under optimal conditions (Kaiser *et al.* 2020). By comparison, increased *g*_s_ has a significant effect on photosynthetic CO_2_ assimilation under FL in *Arabidopsis thaliana* and rice (Kimura *et al.* 2020; Yamori *et al.* 2020). These results indicate that the effects of altered *g*_s_ kinetics on photosynthesis under FL is species dependent. In MN- and LN-plants, the relatively slower kinetics of *g*_s_ led to a higher *L*_gs_ of *A*_N_ during the initial 15 min after transition from low to high light (Fig. 6). Therefore, altered *g*_s_ kinetics would have more significant effects on photosynthetic carbon gain in crop plants grown under low N concentrations.

Many previous studies have indicated that *g*_m_ act as a major limitation for steady-state *A*_N_ under high light in many angiosperms (Peguero-Pina *et al.* 2017; Théroux-Rancourt & Gilbert 2017; Yang, Tong, Yu, Zhang & Huang 2018a; Huang, Yang, Wang & Hu 2019). Increasing *g*_m_ has been thought to be a potential target for improving crop productivity and water use efficiency under constant high light (Flexas *et al.* 2013; Gago *et al.* 2016). However, the limitation of *g*_m_ imposed to *A*_N_ under FL is poorly understood. Upon transition from dark to light, the induction response of *g*_m_ was much faster than that of *g*_s_, leading to the smallest limitation of *g*_m_ imposed to *A*_N_ in *Arabidopsis thaliana* and tobacco (Sakoda, Yamori, Groszmann & Evans 2021). Consequently, one concluded that altering *g*_m_ kinetics would have little impact on *A*_N_ under FL. However, we found that, after transfer from low to high light, *L*_gm_ was higher than *L*_gs_ in tomato plants (Fig. 6). Furthermore, the time to reach 90% of *A*_N_ was closer to that of *g*_m_ rather than that of *g*_s_ (Fig. 3). Therefore, altering *g*_m_ kinetics would significantly influence *A*_N_ upon transfer from low to high light, at least in tomato. These results suggested that the photosynthetic limitation upon transfer from low to high light was largely different from the photosynthetic induction during illumination of dark-adapted leaves. Improving the induction rate of *g*_m_ has a potential to enhance carbon gain and plant biomass under natural FL conditions.

A recent study reported that, if RuBP regeneration limitation was assumed, electron transport imposed the greatest limitation to *A*_N_ during illumination of dark-adapted leaves (Sakoda *et al.* 2021). Based on this result, it is hypothesized that increased activation of electron transport has the potential to enhance carbon gain under naturally FL environments. Controversially, our present study indicated that electron transport was rapidly activated upon transfer from low to high light. After transition from low to high light, the ETR/(*A*_N_ + *R*_d_) value rapidly increased to the peak within 1-2 min and then gradually decreased over time (Fig. 7). These results indicated that, upon transfer from low to high light, the induction response of electron transport was much faster than that of *A*_N_, which was consistent to the photosynthetic performance in rice (Yamori, Makino & Shikanai 2016b). Therefore, induction response of *A*_N_ after transition from low to high light was hardly limited by electron transport in tomato. The effect of electron transport on *A*_N_ upon transition from low to high light is largely different from that upon transition from dark to light. Therefore, to improve photosynthesis under FL in tomato, more attention should be focused on the induction kinetics of CO_2_ diffusional conductance rather than the activation of electron transport.

## Conclusions

We studied the effects of leaf N content on photosynthetic induction after transfer from low to high light in tomato. The induction speeds of *A*_N_, *g*_s_ and *g*_m_ significantly decreased with the decrease in leaf N content. Such delayed photosynthetic induction in plants grown under low N concentration caused a larger loss of carbon gain under FL conditions, which further explained why N deficiency reduced plant biomass under natural FL environments. After transition from low to high light, increasing the induction responses of *g*_s_ and *g*_m_ has the potential to improve *A*_N_ in tomato, especially when plants are grown under low N concentration, whereas photosynthetic induction of *A*_N_ was hardly limited by electron transport. Therefore, altering induction kinetics of CO_2_ diffusional conductance is likely the most effective target for improving photosynthesis under FL conditions in tomato.

## Acknowledgments

This work was supported by the National Natural Science Foundation of China (No. 31971412, 32171505), and the Project for Innovation Team of Yunnan Province.

