## Supporting information for "Photosynthetic induction upon transfer from low to high light is affected by leaf nitrogen content in tomato"

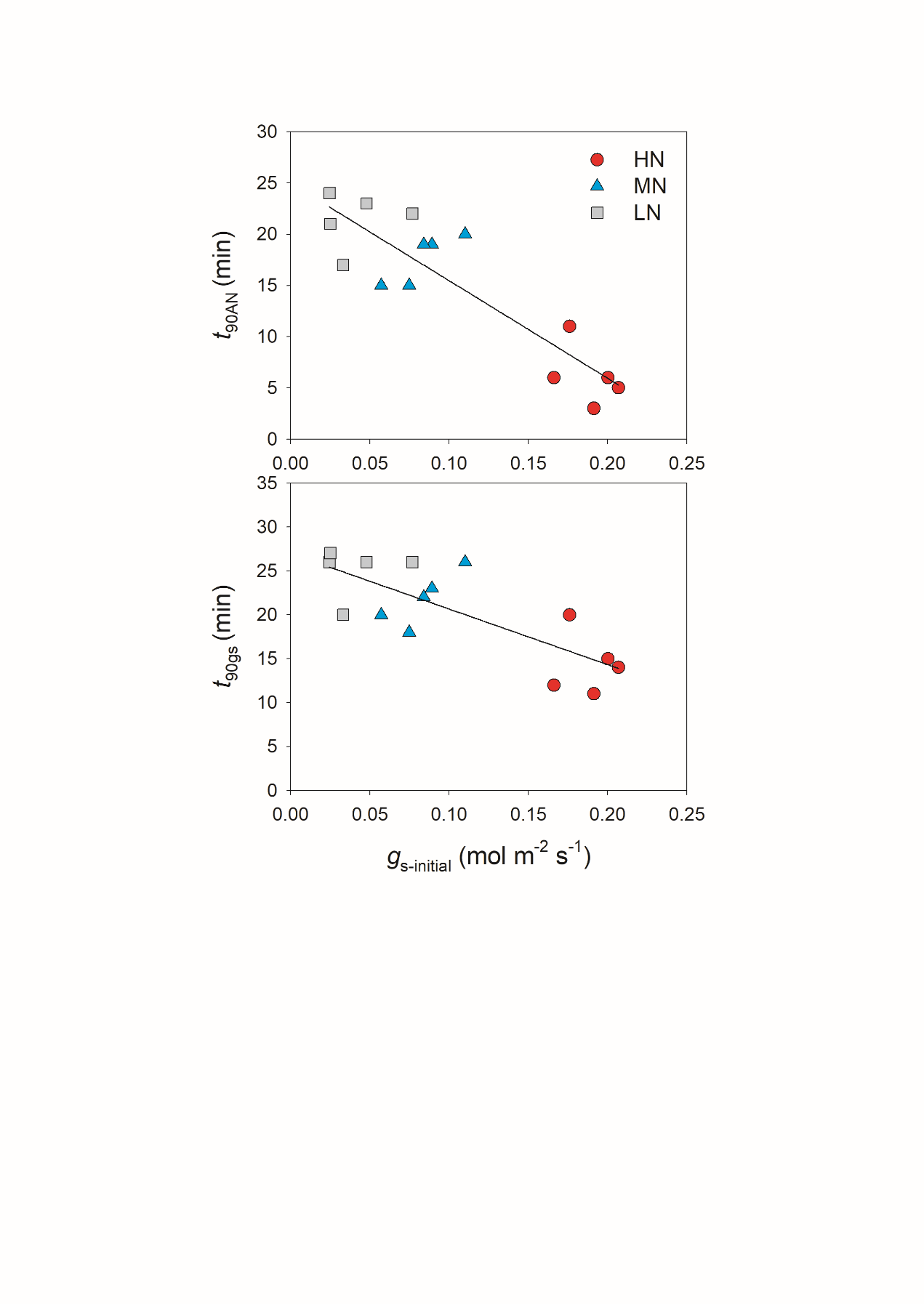

**Figure S1.** Relationships between *t*_90AN_, *t*_90gs_ and the initial *g*_s_ prior to light change. HN, MN and LN represent tomato plants grown under high, medium and low N concentrations, respectively.

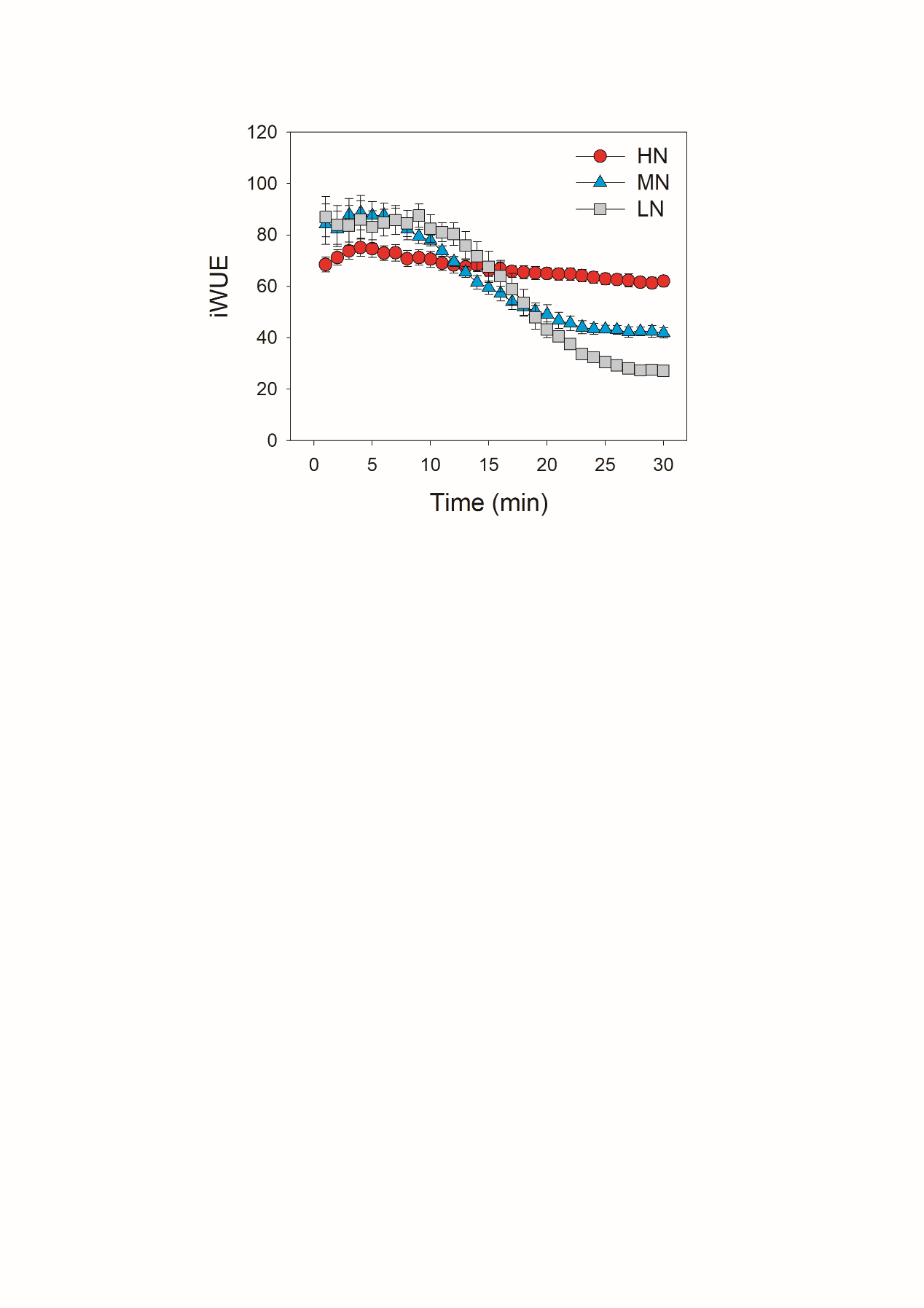

**Figure S2.** Response of intrinsic water use efficiency (iWUE) after transition from 50 to 1500 μmol photons m^−2^ s^−1^. Values are means ± SE (n = 5). HN, MN and LN represent tomato plants grown under high, medium and low N concentrations, respectively.
